# Differential Regulation of Branched-Chain Amino Acids During Early Germination of Mungbean (*Vigna radiata* L.)

**DOI:** 10.64898/2026.07.31.741962

**Authors:** Chanwook Kim, Hakyung Kwon, Sung Don Lim, Yeon-Ji Jo, Jungmin Ha

## Abstract

Branched-chain amino acids (BCAAs) are essential amino acids involved in protein synthesis and energy metabolism. Because animals cannot synthesize BCAA *de novo*, plant-derived BCAAs are important to human nutrition. Although mungbean sprouts are widely consumed as functional plant-based food materials, changes in individual BCAA accumulation and their transcriptional regulation during mungbean germination remain poorly understood. In this study, amino acid contents and transcriptomic profiles were analyzed at three germination stages, 8H, 24H, and 72H. Total BCAA content increased during germination, whereas individual BCAAs exhibited distinct temporal accumulation patterns. Isoleucine and valine increased until 72H, while leucine increased during early germination and decreased after 24H. Transcriptome analysis revealed time-dependent expression changes in BCAA biosynthesis and degradation genes associated with the leucine decrease after 24H. These findings suggest that 24H represents an important transition point for BCAA accumulation and compositional change during mungbean germination. This study provides molecular evidence for the regulation of BCAA metabolism during mungbean germination and supports the potential use of germinated mungbean as a plant-based amino acid resource.

## 1. Introduction

Branched-chain amino acids (BCAAs), including leucine, isoleucine, and valine, are essential amino acids that play key roles in muscle protein synthesis and energy metabolism, contributing to exercise performance and recovery from fatigue in humans (Hoffman & Falvo, 2004; Kaspy et al., 2024; Neinast et al., 2019; Platell et al., 2000; Shimomura et al., 2004). Among the BCAAs, leucine is recognized as a key regulator of protein synthesis, and an increase in its relative proportion has been reported to improve exercise performance and protein metabolism (Duan et al., 2017; Li et al., 2011; Norton & Layman, 2006). Despite their physiological importance, humans and other animals cannot synthesize BCAAs *de novo* and must obtain them through dietary intake (Binder, 2010; Hildebrandt et al., 2015). In this context, plant-derived foods are important sources of dietary BCAAs, and BCAA accumulation in edible plant tissues is closely related to nutritional quality (Sá et al., 2020).

Mungbean (*Vigna radiata* L.) is a major legume crop consumed in both seed and sprout forms. It is considered an important nutritional resource, particularly in developing countries, because of its short growth cycle, low cultivation cost, and adaptability to diverse growing conditions(Hwang et al., 2023; Pataczek et al., 2018). Mungbean is valued for its relatively high protein content and balanced amino acid composition, making it a useful plant-based source of dietary amino acids (Bartholomae et al., 2019; Yi-Shen et al., 2018). In addition to amino acid composition, mungbean is also rich in various functional secondary metabolites, including catechin, chlorogenic acid, and several isoflavonoids, which contribute to its nutritional and health-promoting properties(Jeon et al., 2023; Kim et al., 2023). In particular, mungbean sprouts have attracted increasing attention as functional food material, because germination can enhance the accumulation of bioactive compounds and improve nutritional quality(Chen et al., 2019; Guo et al., 2012; Lim, Kim, et al., 2022).

Germination is a critical developmental transition, which involves the mobilization of stored reserves to support early seedling growth(Nonogaki, 2008). This process requires coordinated regulation of protein synthesis and degradation, energy metabolism, and the redistribution of carbon and nitrogen fluxes across multiple metabolic pathways(Hildebrandt et al., 2015; Zhao et al., 2018). During this process, BCAAs occupy an important metabolic position because they can serve both as structural components for newly synthesized proteins and as metabolic substrates that are catabolized to support energy production(Binder, 2010). Therefore, BCAA accumulation is likely to change dynamically during germination, and understanding these temporal changes is important for the effective utilization of germinated mungbean as a plant-based amino acid resource.

Several key enzymes are known to participate in BCAA biosynthesis and degradation. In BCAA biosynthesis, enzymes such as acetohydroxy acid synthase, ketol-acid reductoisomerase (KARI), dihydroxy-acid dehydratase (DHAD), branched-chain amino acid aminotransferase (BCAT), and 3-isopropylmalate dehydratase (leuC) are involved in the sequential production of leucine, isoleucine, and valine(Binder, 2010; Binder et al., 2007; Xing & Last, 2017). BCAA degradation also involves BCAT family members, together with degradation-associated enzymes such as the branched-chain α-keto acid dehydrogenase complex (BCKDH) and dihydrolipoamide dehydrogenase (DLD), which contribute to the conversion of BCAAs into downstream metabolites that can enter energy metabolism(Binder, 2010; Bo & Fujii, 2024; Hildebrandt et al., 2015; Peng et al., 2015). Therefore, examining the expression patterns of genes encoding these enzymes may help explain temporal changes in individual BCAA accumulation during mungbean germination.

Previous studies on mungbean germination have mainly focused on changes in secondary metabolites, including phenolic compounds, flavonoids, and isoflavones. To the best of our knowledge, few studies have examined individual BCAA dynamics together with the expression of biosynthetic and degradative genes during mungbean germination. This study quantified total amino acid and individual BCAA contents across successive germination stages of mungbean, and analyzed the transcriptome-level expression patterns of genes in the BCAA biosynthesis and degradation pathways. By integrating amino acid profiles with RNA-seq data, this study aimed to identify key germination stages and candidate metabolic genes associated with differential regulation of individual BCAAs.

## 2. Materials and Methods

### 2.1. Plant materials and sample preparation

Mungbean (*Vigna radiata* L.) seeds of the cultivar “Sanpo” were cultivated and harvested in the experimental greenhouse at Gangneung-Wonju National University (Gangneung, Korea; 37.77°N, 128.86°E). Mungbean sprouts were grown according to a previously described method with minor modifications(Kim et al., 2021). Seeds were soaked in distilled water at 37 °C for 16 hours (H) in an incubator chamber (ISS-4075R, JeioTech, Korea) under dark conditions and subsequently germinated in a plant growth chamber (ST001A, Sundotcom, Korea) equipped with an automated spray-irrigation system that supplied water for two minutes every four hours. Sprouts were sampled at 8H, 24H, and 72H after germination (Fig. S1). Fresh tissues were immediately frozen in liquid nitrogen and stored at −80 °C. Portions of each sample were allocated for RNA extraction, whereas the remaining samples were lyophilized (FD 8508, ilShin Biobase, Korea) for five days and finely ground for amino acid analysis. In this manuscript, ungerminated seeds were designated as 0H, whereas germinated samples were denoted according to the time elapsed after germination (8H, 24H, and 72H).

### 2.2. Determination of amino acids and BCAA contents

The contents of amino acids, including BCAA, in mungbean seeds and sprouts were analyzed using an automated amino acid analyzer (L-8800, Hitachi, Japan) at the Center for Research Facilities, Gangneung-Wonju University. Lyophilized samples (50 mg) were hydrolyzed with 10 mL of 6 N HCl in sealed tubes at 110 °C for 22 h. After hydrolysis, the samples were evaporated under reduced pressure, reconstituted in 0.02 N HCl, and filtered through a 0.20-µm syringe filter prior to analysis. Amino acids were separated using an ion-exchange column (4.6 × 60 mm; 2622SC-PH, Hitachi, Japan) at 57 °C, followed by post-column derivatization using a ninhydrin reaction system with the reaction coil set at 135 °C. Gradient elution was performed using buffer solutions PH-1 (05112-79), PH-2 (05113-79), PH-3 (05114-79), PH-4 (05115-79), and RG (05116-79) supplied by Kanto Chemical Co., Inc. (Japan) via Pump 1. Ninhydrin reagent (R1; 299-70501, Wako Pure Chemical Industries, Ltd., Japan), buffer solution (R2; Wako Pure Chemical Industries, Ltd., Japan), and distilled water (R3) were delivered via Pump 2. The detailed gradient program and flow rates are provided in Table S1. The injection volume was set to 20 µL, and eluted amino acids were detected spectrophotometrically at wavelengths ranging from 440 to 570 nm. Quantification was conducted using a certified amino acid standard mixture (Type H[CRM], Wako Pure Chemical Industries, Ltd., Japan). All measurements were performed with three independent biological replicates.

### 2.3. RNA extraction, library construction, and sequencing

Total RNA was extracted from mungbean sprouts using a Ribospin™ Plant RNA extraction Kit (GeneAll, Seoul, Korea) according to the manufacturer’s protocol. For samples collected at 8H, a Ribospin™ Seed/Fruit RNA Extraction Kit (GeneAll, Seoul, Korea) was used. cDNA libraries for RNA-seq were constructed using a TruSeq Stranded mRNA LT Sample Prep Kit (Illumina Inc., CA, USA). A total of nine libraries were prepared from mungbean sprouts sampled at 8H, 24H, and 72H after germination, with three biological replicates per time point. The size and quality of the libraries were evaluated using a 2100 Bioanalyzer (Agilent Technologies, CA, USA). Sequencing was performed in paired-end mode using a TruSeq SBS Kit on the Illumina NovaSeq 6000 platform. The RNA sequencing data were deposited in the NCBI Sequence Read Archive under BioProject accession PRJNA1467411.

### 2.4. Preprocessing, alignment, and identification of DEGs

The mungbean reference genome and gene annotation files were obtained from the Crop Genomics Laboratory, Seoul National University (http://plantgenomics.snu.ac.kr). The reference dataset corresponded to *Vigna radiata* (mungbean) accession VC1973A, assembly version 7, annotation version 1 (2021)(Ha et al., 2021). A total of nine RNA-seq libraries were generated, including three biological replicates per condition. Quality control of raw sequencing reads was performed using FastQC. Raw sequencing reads were aligned to the mungbean reference genome using HISAT2(Kim et al., 2015). SAMtools was used for file format conversion and sorting(Danecek et al., 2021). Gene-level read counts were quantified using featureCounts(Liao et al., 2014). Count data were normalized using the trimmed mean of M-values (TMM) method, and differential expression analysis was performed using edgeR(Robinson et al., 2010). Differentially expressed genes (DEGs) were defined as genes with an absolute log₂ fold change (|log₂FC|) ≥ 1 and a false discovery rate (FDR) < 0.05.

### 2.5. Functional annotation and time-course expression analysis

Functional enrichment analyses for Gene Ontology (GO) terms and The Kyoto Encyclopedia of Genes and Genomes (KEGG) pathways were performed using the clusterProfiler package(Consortium, 2004; Yu et al., 2012). To group genes into temporal expression clusters, time-course expression patterns were analyzed with TCseq(Mengjun, 2019), while TPM values were transformed to Z-scores. For pathway-level visualization, gene expression values were represented as log_2_(CPM + 1), while differential changes between consecutive stages were shown using log₂FC values.

### 2.6 Statistical analysis

Statistical analyses were performed using R(Team, 2016). Significant differences were tested using one-way analysis of variance (ANOVA), followed by Duncan’s multiple range test. *p* value < 0.05 was defined as statistical significance, and the data are presented as means ± SD.

## 3. Results

### 3.1. Changes in amino acid and BCAA contents during mungbean germination

Changes in total amino acid and BCAA contents were analyzed during mungbean germination (Fig. 1). Total amino acid content increased progressively from seed stage (0H, 192.95 ± 0.89 mg/g) to significantly higher levels at 24H (246.54 ± 5.20 mg/g) and 72H (252.17 ± 2.93 mg/g) (Fig. 1a). BCAA content also shifted over germination time, showing a gradual increase from 8H (25.83 ± 0.52 mg/g) onward compared with 0H (23.49 ± 0.89 mg/g). At 24H (31.33 ± 1.18 mg/g) and 72H (36.82 ± 0.59 mg/g), BCAA levels were significantly elevated, indicating progressive accumulation of BCAA during germination. Analysis of individual BCAA components—leucine (Leu), isoleucine (Ile), and valine (Val)—revealed distinct temporal patterns for each amino acid (Fig. 1b). Leu content increased from 3.85 ± 0.20 mg/g at 0H to a maximum at 24H (16.84 ± 0.70 mg/g), followed by a decrease at 72H (15.45 ± 0.31 mg/g). In contrast, Ile and Val exhibited continuous increases throughout germination, reaching their highest levels at 72H (Ile, 8.85 ± 0.13 mg/g; Val, 12.52 ± 0.14 mg/g) compared with 0H (Ile, 5.92 ± 0.36 mg/g; Val, 5.63 ± 0.23 mg/g). Overall, total BCAA content increased during mungbean germination. Leucine decreased after 24H, while isoleucine and valine exhibited sustained accumulation.

**Fig. 1.**
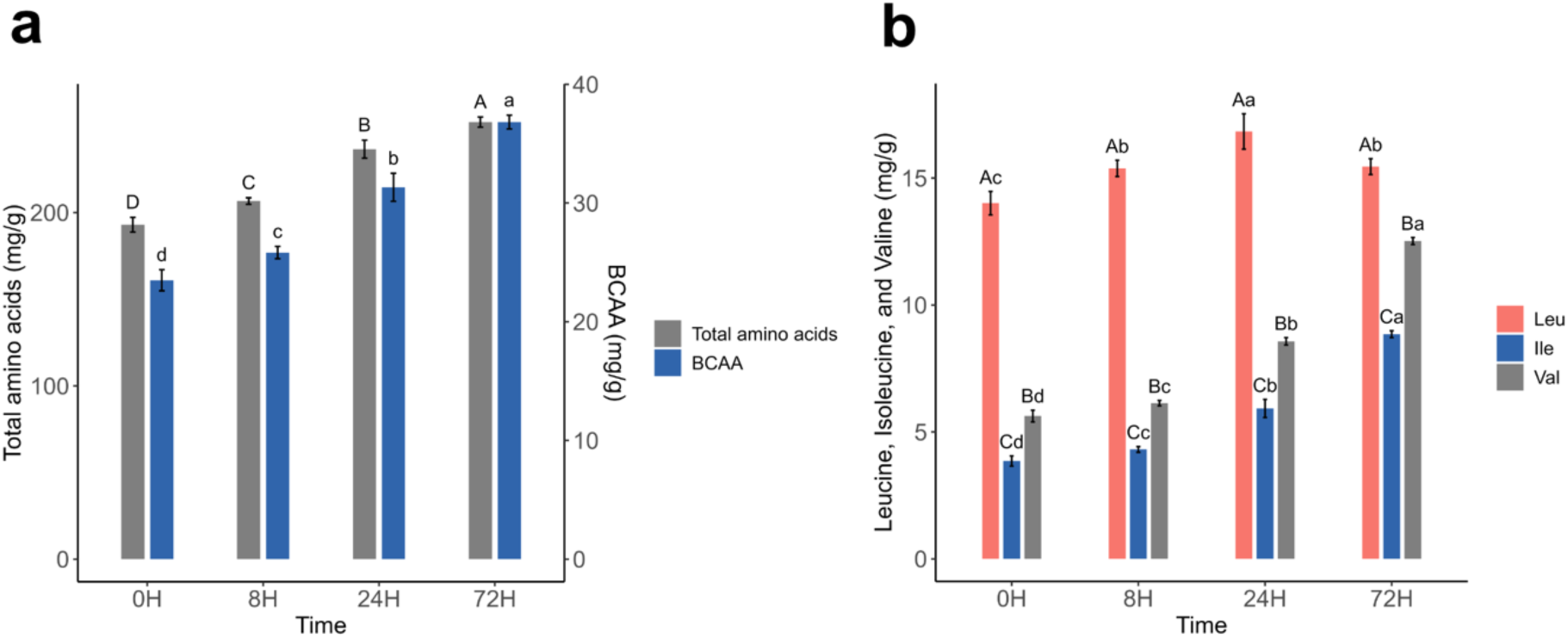
Changes in total amino acid and BCAA contents during mungbean germination. (a) Total amino acids and BCAA content were measured at 0H (seed), 8H, 24H, and 72H after germination. (b) Changes in individual BCAA components: leucine (Leu), isoleucine (Ile), and valine (Val). Data are presented as means ± SD (n = 3). Different letters indicate significant differences among time points according to Duncan’s multiple range test (*p* < 0.05).

### 3.2. Time-course RNA-seq profiling of mungbean germination

RNA-seq analysis was conducted to examine transcriptomic changes during mungbean germination at 8H, 24H, and 72H. A total of 5.1–7.7 Gb of sequence data was generated per library, corresponding to 50.52–76.25 million reads per sample (Table S2). The GC content ranged from 44.8% to 45.5% across all samples, and base quality scores were consistently high, with Q20 and Q30 values of approximately 97.7–98.0% and 93.6–94.3%, respectively. Alignment of reads to the mungbean reference genome showed high mapping rates ranging from 97.2% to 97.5%. Differential expression analysis was performed by comparing transcript levels between consecutive germination stages. In the comparison between 8H and 24H, a total of 11,606 DEGs were identified, including 7,057 upregulated and 4,549 downregulated genes (Fig. S2a). In the comparison between 24H and 72H, 7,563 DEGs were detected, of which 4,071 genes were upregulated and 3,492 genes were downregulated. In total, 4,583 DEGs were commonly identified in both the 8H–24H and 24H–72H comparisons (Fig. S2b). While 7,023 DEGs were uniquely detected in the 8H–24H comparison, 2,980 DEGs were specific to the 24H–72H comparison. The substantial number of DEGs detected at both intervals indicates that mungbean germination is accompanied by extensive transcriptomic reprogramming, and the marked difference in DEG counts between the two intervals further reflects the highly dynamic nature of this developmental transition.

### 3.3. Functional enrichment analysis of DEGs by timepoint

To characterize transcriptomic changes during mungbean germination, Gene Ontology (GO) and Kyoto Encyclopedia of Genes and Genomes (KEGG) pathway enrichment analyses were performed using DEGs identified from the 8H–24H and 24H–72H comparisons (Fig. 2). GO enrichment analysis showed that DEGs from the 8H–24H comparison were primarily associated with transport and carbohydrate metabolism, whereas those from the 24H–72H comparison were enriched in functions related to nucleic acid metabolism and protein complex formation (Fig. 2a, b). KEGG pathway enrichment analysis revealed several pathways commonly enriched in both comparisons, including “plant hormone signal transduction”, “starch and sucrose metabolism”, and “phenylpropanoid biosynthesis” (Fig. 2c, d). In the 8H– 24H comparison, DEGs were mainly enriched in pathways related to energy production, such as “starch and sucrose metabolism” and “glycolysis/gluconeogenesis” (Fig. 2c). In contrast, the 24H–72H comparison showed enrichment of pathways associated with signaling and secondary metabolism, including “MAPK signaling pathway”, “plant–pathogen interaction”, and “flavonoid biosynthesis” (Fig. 2d). These results indicate that mungbean germination involves a temporal shift in transcriptional programs, from carbohydrate mobilization and energy production during early germination (8H–24H) to signaling and secondary metabolism at later stages (24H–72H).

**Fig. 2.**
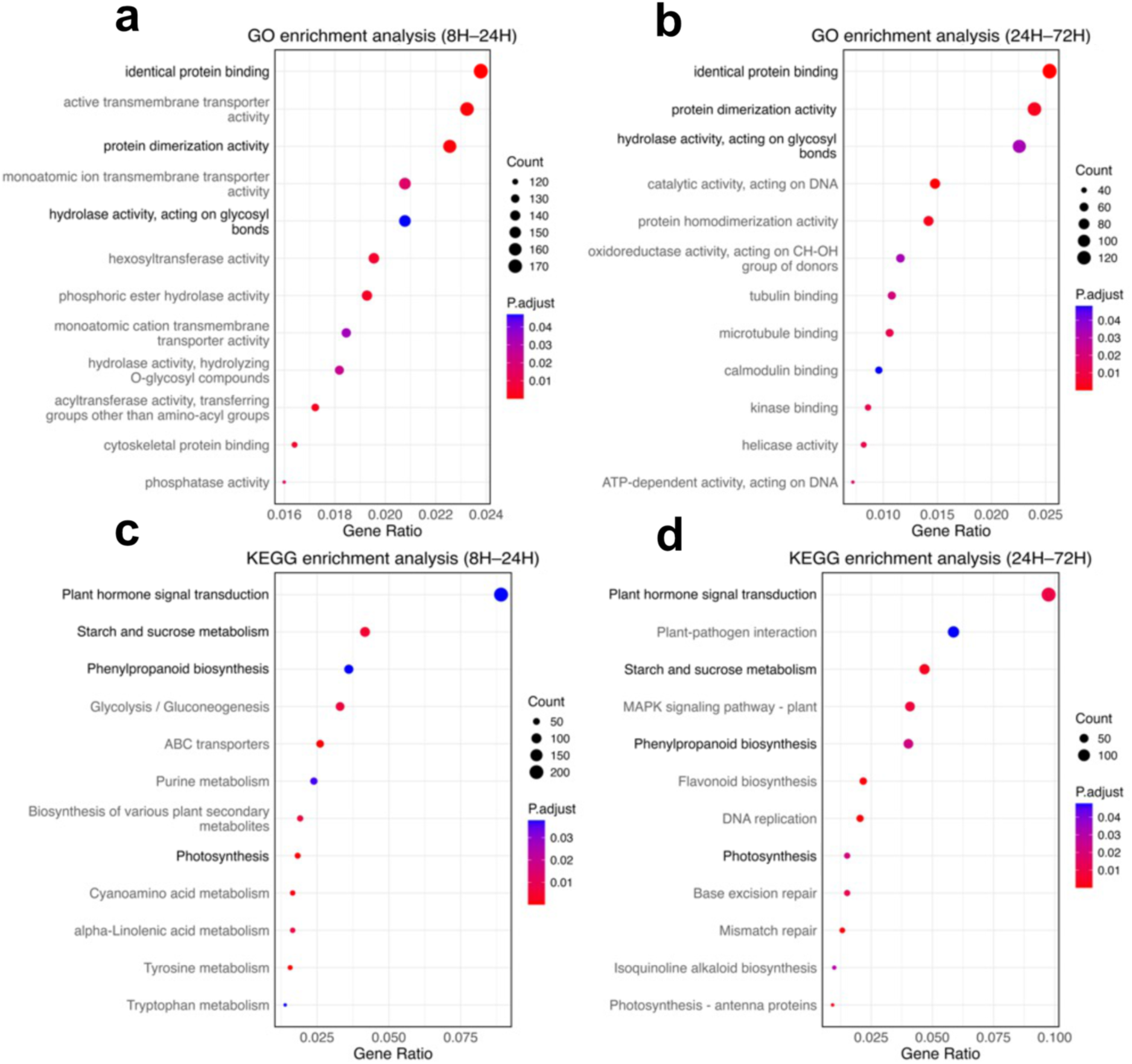
Gene Ontology (GO) and Kyoto Encyclopedia of Genes and Genomes (KEGG) enrichment analysis of DEGs during mungbean germination. (a, b) GO enrichment results for DEGs in the 8H–24H (a) and 24H–72H (b) comparisons. (c, d) KEGG pathway enrichment results for DEGs in the 8H–24H (c) and 24H–72H (d) comparisons. Circle size represents the number of genes, and color indicates adjusted *p* values (P.adjust). Terms and pathways commonly enriched in both comparisons are highlighted.

### 3.4. Time-series clustering of DEGs during mungbean germination

To analyze time-dependent gene expression patterns during mungbean germination, time-series clustering analysis was performed using 4,583 DEGs that were commonly identified both in the 8H–24H and 24H–72H comparisons. Based on their expression patterns, these commonly identified DEGs were classified into four distinct clusters, each exhibiting a unique expression trend across germination stages (Fig. 3). Clusters 1 to 4, comprising 1,916, 1,467, 631, and 569 genes, respectively, were then subjected to GO enrichment analysis to investigate their biological roles (Fig. S3). Cluster 1 showed low expression levels at 8H followed by a gradual increase, with the highest expression observed at 72H. This cluster was predominantly associated with energy-related processes, including “generation of precursor metabolites and energy” and “photosynthesis,” as well as stress response terms such as “response to water deprivation.” Cluster 2, which displayed increased expression between 8H and 24H, prior to a decline at 72H, was enriched in cell cycle–related processes, including “mitotic cell cycle” and “nuclear division”. Cluster 3, which exhibited relatively low expression at 24H and increased expression observed at 72H, was primarily associated with responses to oxygen availability, including “response to hypoxia” and “response to decreased oxygen levels”. Lastly, cluster 4, exhibiting a continuous decrease in expression as germination progressed, was enriched in RNA processing–related terms, including “RNA modification” and “ribosome biogenesis”. Genes involved in the BCAA metabolic pathway were primarily found in Clusters 1 and 2. BCAT (Vradi02g00002555.1) and leuC (chloroplastic 3-isopropylmalate dehydratase large subunit Vradi07g00000549.1) were assigned to Cluster 1, showing an overall upward expression pattern during germination. In contrast, DHAD (Vradi08g00000722.1) and DLD (chloroplastic dihydrolipoyl dehydrogenase 1, Vradi02g00004047.1) belonged to Cluster 2 and exhibited transient up-regulation at 24H followed by decreased expression at 72H.

**Fig. 3.**
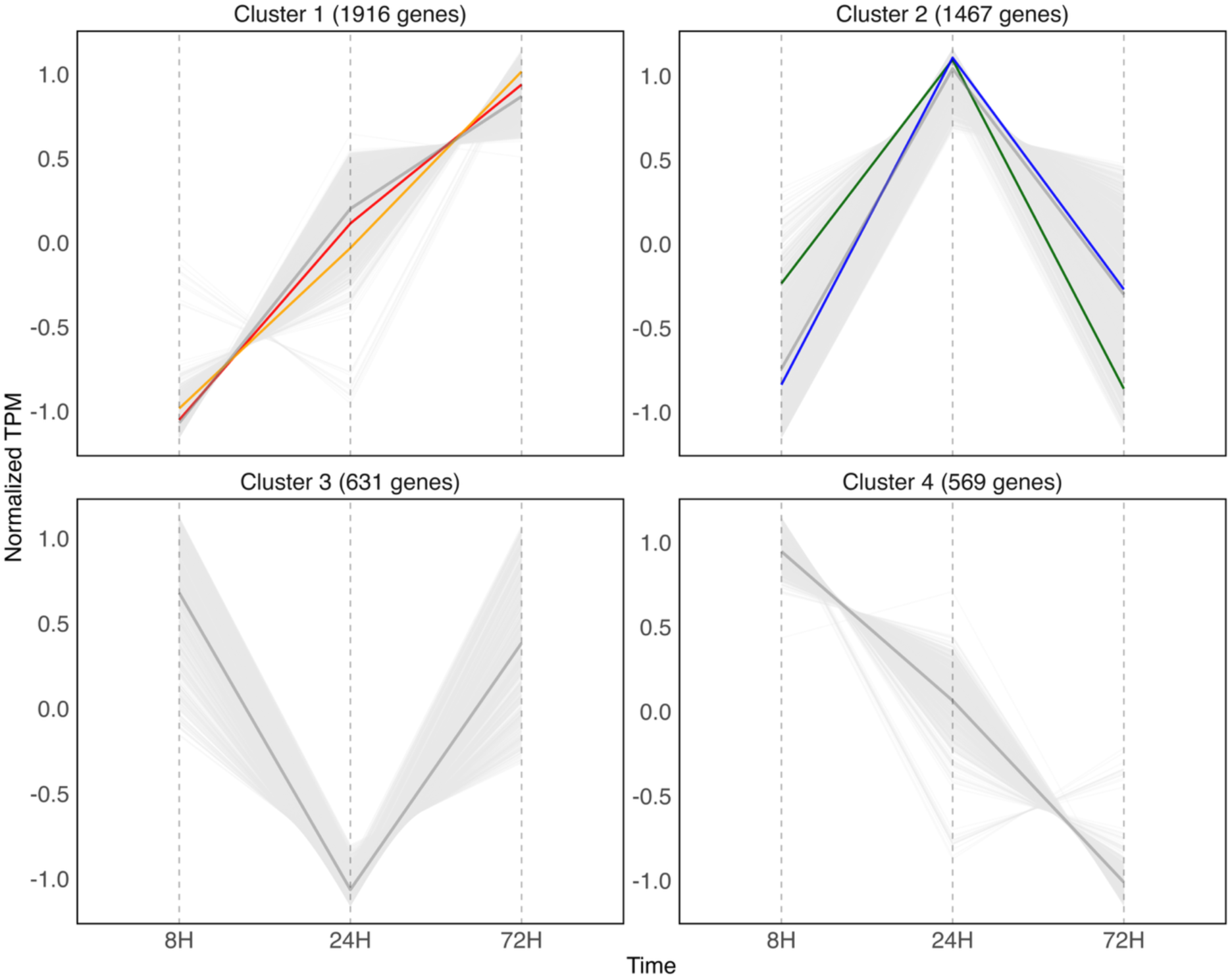
Time-course expression patterns of DEGs commonly identified in both the 8H–24H and 24H–72H comparisons were analyzed using the TCseq package. A total of 4,583 shared DEGs were grouped into four distinct clusters based on their normalized transcripts per million (TPM, Z-score) expression profiles. The gray central line in each panel represents the average expression profile of all genes assigned to the corresponding cluster, whereas light gray lines indicate individual gene expression trajectories within each cluster. Colored lines highlight the expression patterns of representative genes associated with BCAA metabolism: BCAT (red line, Vradi02g00002555.1), leuC, chloroplastic 3-isopropylmalate dehydratase large subunit (orange line, Vradi07g00000549.1), DHAD, dihydroxy-acid dehydratase (green line, Vradi08g00000722.1), and DLD, chloroplastic dihydrolipoyl dehydrogenase 1 (blue line, Vradi02g00004047.1). The number of genes assigned to each cluster is indicated at the top of each panel. Gene identifiers shown in the pathway diagram are based on KEGG Orthology (KO) annotations.

### 3.5 BCAA biosynthesis and degradation pathways during mungbean germination

To investigate the leucine-specific decrease observed after 24H of mungbean germination, genes involved in BCAA biosynthesis and degradation pathways were identified, and their transcriptional profiles were compared using log_2_(CPM + 1) values at 8H, 24H, and 72H (Fig. 4). Several genes associated with leucine biosynthesis showed expression patterns that were partially consistent with leucine accumulation, exhibiting relatively high expression at 24H followed by decreased expression at 72H. For example, DLD (Vradi02g00004047.1) showed expression values of 4.48, 6.26, and 5.09 at 8H, 24H, and 72H, respectively, and leuB (3-isopropylmalate dehydrogenase; Vradi10g00001380.1) showed values of 5.09, 6.36, and 5.95. Similar decreasing trends after 24H were also observed in some PDHA-, PDHB-, leuA-, and leuC-related genes. In contrast, several genes involved in leucine degradation maintained or increased their expression at later germination stages. Among ACAT (acetyl-CoA C-acetyltransferase) paralogues, Vradi09g00000859.1 increased continuously during germination, with expression values of 7.32, 7.92, and 8.39 at 8H, 24H, and 72H, respectively, whereas Vradi06g00000825.1 showed increased expression after 24H, with values of 8.38, 7.21, and 7.75, respectively. These results suggest that reduced expression of several leucine biosynthesis-related genes after 24H, together with the maintenance or increase of leucine degradation-related genes, may be associated with reduced leucine accumulation at later germination stages.

**Fig. 4.**
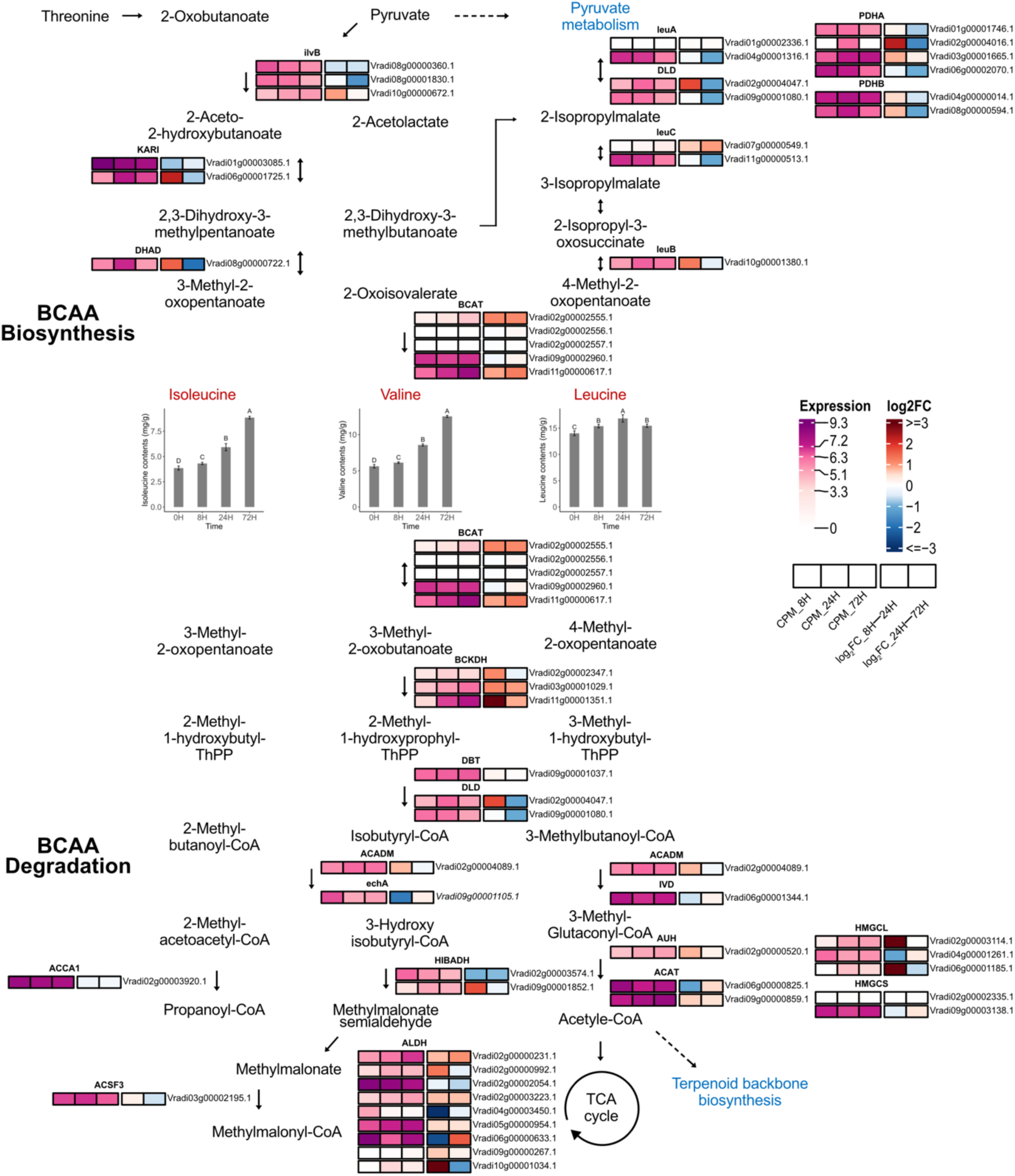
Changes in gene expression associated with BCAA biosynthesis and degradation pathways during mungbean germination. The biosynthesis and degradation pathways of isoleucine, valine, and leucine were reconstructed based on KEGG pathway annotations and mungbean gene annotation information to illustrate BCAA metabolism during mungbean germination. Enzyme names are shown above the heatmap boxes, and the corresponding mungbean gene IDs assigned to each enzyme are listed to the right of the boxes. For each gene, the first three heatmap boxes represent expression levels at 8H, 24H, and 72H based on log_2_(CPM + 1) values. In the expression heatmap, colors closer to white indicate lower expression levels, whereas colors closer to red indicate higher expression levels. The last two boxes represent sequential expression changes based on log₂ fold change values for 8H–24H and 24H–72H. In the log_2_FC heatmap, red indicates up-regulation, blue indicates down-regulation, and white indicates little or no change.

Although several paralogues described above showed expression patterns consistent with changes in amino acid contents, not all paralogues of BCAA metabolism genes followed this pattern. Among the two leuC paralogues, Vradi07g00000549.1 showed a continuous increase in expression during germination. Similarly, among the five BCAT paralogues, Vradi02g00002555.1 showed a continuous increase in expression during germination, with expression values of 1.79, 2.93, and 4.02 at 8H, 24H, and 72H, respectively. Another BCAT paralogue, Vradi02g00002557.1, remained at low expression levels. In addition, the BCKDH paralogue Vradi11g00001351.1 showed increased expression at later germination stages, with values of 3.55, 6.82, and 7.65. These results suggest that only a subset of BCAA metabolism-related paralogues showed expression patterns associated with changes in amino acid contents, whereas other paralogues annotated with the same enzymatic functions exhibited distinct temporal profiles during germination.

## Discussion

With the growing interest in plant-based diets, plant-derived protein sources have received increasing attention as alternatives or complements to animal-derived proteins(Andreani et al., 2023; Sá et al., 2020; Semba et al., 2021). Among legumes, soybean is one of the most extensively studied plant-based protein sources, whereas the nutritional value of mungbean, particularly its potential as a source of BCAA, has received comparatively limited attention(Sui et al., 2021; Tang et al., 2014). In the present study, total BCAA content in mungbean increased from 23.49 mg/g in seeds to a maximum of 36.82 mg/g during germination (Fig. 1). Notably, this value exceeded those reported for several commonly consumed soybean-derived foods, including soy drink (5.72 mg/g), soy dessert (5.20 mg/g), and tofu curd (25.08 mg/g)(Kudełka et al., 2021). Given that mungbean is predominantly consumed as sprouts, these findings suggest that its nutritional value as a dietary source of BCAAs may have been underestimated(Gan et al., 2017). Therefore, mungbean has nutritional value as an amino acid source, including BCAAs, and may serve as a complementary plant-based resource to soybean-centered plant protein materials.

Although total BCAA content increased during germination, the three constituent amino acids exhibited distinct accumulation patterns (Fig. 1). Leucine content decreased after 24H of germination, whereas isoleucine and valine continued to accumulate, consistent with previous observations in cultivated and wild mungbean germplasm(Kim et al., 2025). This finding is nutritionally important because BCAA supplements are commonly formulated with leucine, isoleucine, and valine at a ratio of 2:1:1, and a relatively high leucine proportion has been associated with enhanced muscle protein synthesis and exercise performance(Arroyo-Cerezo et al., 2021; Osmond et al., 2019). Therefore, evaluation of germinated mungbean as a dietary BCAA source should consider not only total BCAA content but also changes in individual BCAA composition during germination. From this perspective, the 24H germination stage appears to provide the most favorable balance between increased total BCAA content and maintenance of a relatively high leucine proportion(Kim et al., 2025).

The decline in leucine content after 24H of germination may reflect a metabolic shift toward increased leucine utilization during dark germination. Because seedlings germinated in the absence of light rely primarily on the mobilization of stored seed reserves rather than photosynthetic carbon assimilation, amino acids can serve as alternative substrates for energy production(Nonogaki, 2008; Zhao et al., 2018). In particular, leucine can be degraded to acetyl-CoA, thereby contributing to central carbon metabolism(Anderson et al., 1998; Hildebrandt et al., 2015; Latimer et al., 2018). Consistent with this interpretation, Cluster 1 genes, which showed progressively increased expression during germination, were enriched in GO terms related to the “generation of precursor metabolites and energy” as well as “monocarboxylic acid metabolic processes” (Fig. 3, Fig. S3). Such enrichment is consistent with the metabolic reprogramming that occurs during seed germination, a process characterized by high respiratory demand and dynamic adjustment to oxygen availability(Rolletschek et al., 2025). Furthermore, genes involved in leucine biosynthesis showed relatively high expression at early germination stages followed by decreased expression at later stages, whereas genes associated with leucine catabolism and downstream carbon metabolic pathways exhibited sustained or increased expression during germination (Fig. 4). These contrasting expression patterns suggest a coordinated shift from leucine biosynthesis toward metabolic utilization, which may contribute to the observed reduction in leucine content during germination(Heinemann & Hildebrandt, 2021; Peng et al., 2015). Together, these findings suggest that the reduction in leucine content during later stages of germination may result from both decreased biosynthetic activity and increased metabolic utilization of leucine.

Although enzymes commonly involved in the biosynthesis of three BCAAs, including ilvB, KARI, and DHAD, showed reduced or limited expression changes, isoleucine and valine contents continued to increase during germination (Fig. 4). One possible explanation for this observation is that the reduced accumulation of leucine may be associated with altered allocation of shared metabolic precursors within the BCAA pathway. The decrease in leucine content after 24H coincided with reduced expression of leucine biosynthesis-related genes together with increased expression of leucine catabolic processes, suggesting that leucine is more actively utilized during germination. This shift may reduce competition for shared precursors, thereby allowing continued accumulation of isoleucine and valine despite reduced expression of common biosynthetic enzymes(Chen et al., 2010). Overall, these results suggest that carbon flux within the BCAA pathway is differentially allocated among the three amino acids under conditions of reduced biosynthetic activity, with leucine acting as a major metabolic sink that influences the relative accumulation of isoleucine and valine during germination(Heinemann & Hildebrandt, 2021; Peng et al., 2015; Taylor et al., 2004).

The observed leucine-specific decrease after 24H was accompanied by expression changes in only a subset of BCAA metabolism-related paralogues, whereas other paralogues annotated to the same pathway showed distinct temporal profiles during germination (Fig. 4). This suggests that pathway annotation alone may not distinguish paralogues that are functionally responsive under specific physiological conditions. In mungbean, where multiple paralogous genes are present, sequence similarity-based annotation may group genes into the same pathway even though their roles can diverge depending on developmental or metabolic context(Harikrishnan et al., 2015; Panchy et al., 2016). For example, several annotated genes such as Vradi01g00002336.1 and Vradi02g00002556.1 showed low expression levels or no clear temporal variation during germination, indicating limited responsiveness to germination-associated metabolic changes. Therefore, the present results provide transcriptome-based evidence that may help highlight genes showing germination-responsive expression patterns within the annotated BCAA pathway, offering functional insights into pathway-level regulation in mungbean.

BCAT functions in both the final transamination step of BCAA biosynthesis and the initial step of BCAA degradation, and exists as a multigene family with isoform-specific functions(Binder, 2010; Campbell et al., 2001). In plants such as Arabidopsis, chloroplast-and mitochondria-localized BCAT isoforms have been reported to be functionally differentiated, suggesting compartment-specific roles in BCAA metabolism(Diebold et al., 2002; Maloney et al., 2010). In the mungbean genome, five BCAT genes were identified and showed distinct expression levels and temporal patterns during germination (Fig. 4). Based on KEGG annotation information, the five mungbean BCAT genes were annotated as different BCAT types, including chloroplastic BCATs (Vradi02g00002556.1, Vradi09g00002960.1, and Vradi11g00000617.1), a mitochondrial BCAT (Vradi02g00002557.1), and a putative BCAT (Vradi02g00002555.1), suggesting possible functional divergence within the gene family.

Notably, Vradi02g00002555.1, which is currently annotated as a putative BCAT, showed clear differential expression during germination and was consistently identified as a DEG in previous transcriptomic studies in mungbean sprouts(Lim, Kang, et al., 2022). Although its functional characterization remains limited, these results suggest that Vradi02g00002555.1 may play a more important role in BCAA metabolic regulation during germination than its current annotation implies.

This study demonstrates that amino acid composition in mungbean is not simply increased or decreased during germination, but is dynamically reshaped over germination time, accompanied by transcriptional changes in amino acid metabolism-related genes. In particular, although total BCAA content increased during germination, leucine, isoleucine, and valine showed distinct temporal patterns, suggesting that mungbean sprouts with a desired BCAA composition could be obtained by adjusting the germination stage. From this perspective, the 24H germination stage can be considered an important time point at which both increased total BCAA content and leucine composition should be taken into account. In addition, by integrating amino acid quantification with transcriptomic information, this study suggests that changes in BCAA composition during germination may be associated with time-dependent expression changes in related metabolic genes. The expression profiles of genes involved in BCAA biosynthesis and degradation provide basic clues for understanding the molecular background underlying changes in amino acid composition during mungbean germination. Overall, this study shows that the BCAA composition of mungbean sprouts can vary depending on the germination stage and provides fundamental information for the use of germinated mungbean as a functional plant-based amino acid material.

## CRediT authorship contribution statement

**Chanwook Kim:** Resources; Data curation; Software; Formal analysis; Validation; Visualization; Methodology; Writing – original draft.

**Hakyung Kwon:** Data curation; Software; Formal analysis; Investigation; Validation; Visualization; Methodology.

**Sung Don Lim**: Data curation; Validation; Visualization.

**Yeon-Ji Jo:** Conceptualization; Data curation; Software; Visualization; Funding acquisition; Methodology; Writing – review and editing.

**Jungmin Ha:** Conceptualization; Resources; Supervision; Funding acquisition; Methodology; Writing – review and editing; Project administration.

## Acknowledgements

This work was supported by the New Faculty Startup Fund from Seoul National University, the National Research Foundation of Korea (NRF) grant funded by the Korea government (MSIT) (RS-2026-25489549), and the ANCHOR program through the Gangwon ANCHOR Center, funded by the Ministry of Education (MOE) and the Gangwon State (G.S.), Republic of Korea (2026-ANCHOR-10-005).

## Data Availability

The RNA-seq data generated in this study have been deposited in the NCBI Sequence Read Archive under BioProject accession PRJNA1467411. All other data generated or analyzed during this study are included in this article, its supplementary information files, and the accompanying data file containing processed gene expression values.

## Corresponding authors

Correspondence to Jungmin Ha and Yeon-Ji Jo.

## References

1. Anderson, M. D., Che, P., Song, J., Nikolau, B. J., & Wurtele, E. S. (1998). 3-Methylcrotonyl-coenzyme A carboxylase is a component of the mitochondrial leucine catabolic pathway in plants. Plant Physiology, 118(4), 1127–1138.

2. Andreani, G., Sogari, G., Marti, A., Froldi, F., Dagevos, H., & Martini, D. (2023). Plant-based meat alternatives: technological, nutritional, environmental, market, and social challenges and opportunities. Nutrients, 15(2), 452.

3. Arroyo-Cerezo, A., Cerrillo, I., Ortega, A., & Fernandez-Pachon, M.-S. (2021). Intake of branched chain amino acids favors post-exercise muscle recovery and may improve muscle function: optimal dosage regimens and consumption conditions. The Journal of Sports Medicine and Physical Fitness, 61(11), 1478–1489.

4. Bartholomae, E., Incollingo, A., Vizcaino, M., Wharton, C., & Johnston, C. S. (2019). Mung bean protein supplement improves muscular strength in healthy, underactive vegetarian adults. Nutrients, 11(10), 2423.

5. Binder, S. (2010). Branched-chain amino acid metabolism in Arabidopsis thaliana. The Arabidopsis Book/American Society of Plant Biologists, 8, e0137.

6. Binder, S., Knill, T., & Schuster, J. (2007). Branched-chain amino acid metabolism in higher plants. Physiologia Plantarum, 129(1), 68–78.

7. Bo, T., & Fujii, J. (2024). Primary roles of branched chain amino acids (BCAAs) and their metabolism in physiology and metabolic disorders. Molecules, 30(1), 56.

8. Campbell, M. A., Patel, J. K., Meyers, J. L., Myrick, L. C., & Gustin, J. L. (2001). Genes encoding for branched-chain amino acid aminotransferase are differentially expressed in plants. Plant Physiology and Biochemistry, 39(10), 855–860.

9. Chen, H., Saksa, K., Zhao, F., Qiu, J., & Xiong, L. (2010). Genetic analysis of pathway regulation for enhancing branched-chain amino acid biosynthesis in plants. The Plant Journal, 63(4), 573–583.

10. Chen, L., Wu, J. e., Li, Z., Liu, Q., Zhao, X., & Yang, H. (2019). Metabolomic analysis of energy regulated germination and sprouting of organic mung bean (Vigna radiata) using NMR spectroscopy. Food Chemistry, 286, 87–97.

11. Consortium, G. O. (2004). The Gene Ontology (GO) database and informatics resource. Nucleic acids research, 32(suppl_1), D258-D261.

12. Danecek, P., Bonfield, J. K., Liddle, J., Marshall, J., Ohan, V., Pollard, M. O., Whitwham, A., Keane, T., McCarthy, S. A., & Davies, R. M. (2021). Twelve years of SAMtools and BCFtools. Gigascience, 10(2), giab008.

13. Diebold, R., Schuster, J., Däschner, K., & Binder, S. (2002). The branched-chain amino acid transaminase gene family in Arabidopsis encodes plastid and mitochondrial proteins. Plant Physiology, 129(2), 540–550.

14. Duan, Y., Zeng, L., Li, F., Wang, W., Li, Y., Guo, Q., Ji, Y., Tan, B. e., & Yin, Y. (2017). Effect of branched-chain amino acid ratio on the proliferation, differentiation, and expression levels of key regulators involved in protein metabolism of myocytes. Nutrition, 36, 8–16.

15. Gan, R.-Y., Lui, W.-Y., Wu, K., Chan, C.-L., Dai, S.-H., Sui, Z.-Q., & Corke, H. (2017). Bioactive compounds and bioactivities of germinated edible seeds and sprouts: An updated review. Trends in Food Science & Technology, 59, 1–14.

16. Guo, X., Li, T., Tang, K., & Liu, R. H. (2012). Effect of germination on phytochemical profiles and antioxidant activity of mung bean sprouts (Vigna radiata). Journal of agricultural and food chemistry, 60(44), 11050–11055.

17. Ha, J., Satyawan, D., Jeong, H., Lee, E., Cho, K. H., Kim, M. Y., & Lee, S. H. (2021). A near-complete genome sequence of mungbean (Vigna radiata L.) provides key insights into the modern breeding program. The plant genome, 14(3), e20121.

18. Harikrishnan, S. L., Pucholt, P., & Berlin, S. (2015). Sequence and gene expression evolution of paralogous genes in willows. Scientific Reports, 5(1), 18662.

19. Heinemann, B., & Hildebrandt, T. M. (2021). The role of amino acid metabolism in signaling and metabolic adaptation to stress-induced energy deficiency in plants. Journal of Experimental Botany, 72(13), 4634–4645.

20. Hildebrandt, T. M., Nesi, A. N., Araújo, W. L., & Braun, H.-P. (2015). Amino acid catabolism in plants. Molecular plant, 8(11), 1563–1579.

21. Hoffman, J. R., & Falvo, M. J. (2004). Protein–which is best? Journal of sports science & medicine, 3(3), 118.

22. Hwang, N., Gu, B.-J., & Ryu, G.-H. (2023). Physicochemical properties of low-moisture extruded meat analog by replacing isolated soy with mung bean protein.

23. Jeon, S., Kim, B. C., & Ha, J. (2023). Tissue-Specific Metabolic Profiling of Mungbean (Vigna radiata L.) Genotypes with Different Seed Coat Colors. Journal of Food Quality, 2023(1), 7555915.

24. Kaspy, M. S., Hannaian, S. J., Bell, Z. W., & Churchward-Venne, T. A. (2024). The effects of branched-chain amino acids on muscle protein synthesis, muscle protein breakdown and associated molecular signalling responses in humans: an update. Nutrition Research Reviews, 37(2), 273–286.

25. Kim, B. C., Lim, I., & Ha, J. (2023). Metabolic profiling and expression analysis of key genetic factors in the biosynthetic pathways of antioxidant metabolites in mungbean sprouts. Frontiers in Plant Science, 14, 1207940.

26. Kim, B. C., Lim, I., Jeon, S. Y., Kang, M., & Ha, J. (2021). Effects of irrigation conditions on development of mungbean (Vigna radiata l.) sprouts. Plant breeding and biotechnology, 9(4), 310–317.

27. Kim, C., Jeon, S., Jo, Y.-J., & Ha, J. (2025). Amino acids and BCAA composition of Mungbean (Vigna radiata L.) seeds and sprouts for plant-based protein applications. Scientific Reports, 15(1), 29590.

28. Kim, D., Langmead, B., & Salzberg, S. L. (2015). HISAT: a fast spliced aligner with low memory requirements. Nature methods, 12(4), 357–360.

29. Kudełka, W., Kowalska, M., & Popis, M. (2021). Quality of soybean products in terms of essential amino acids composition. Molecules, 26(16), 5071.

30. Latimer, S., Li, Y., Nguyen, T. T., Soubeyrand, E., Fatihi, A., Elowsky, C. G., Block, A., Pichersky, E., & Basset, G. J. (2018). Metabolic reconstructions identify plant 3-methylglutaconyl-CoA hydratase that is crucial for branched-chain amino acid catabolism in mitochondria. The Plant Journal, 95(2), 358–370.

31. Li, F., Yin, Y., Tan, B., Kong, X., & Wu, G. (2011). Leucine nutrition in animals and humans: mTOR signaling and beyond. Amino Acids, 41(5), 1185–1193.

32. Liao, Y., Smyth, G. K., & Shi, W. (2014). featureCounts: an efficient general purpose program for assigning sequence reads to genomic features. Bioinformatics, 30(7), 923–930.

33. Lim, I., Kang, M., Kim, B. C., & Ha, J. (2022). Metabolomic and transcriptomic changes in mungbean (Vigna radiata (L.) R. Wilczek) sprouts under salinity stress. Frontiers in Plant Science, 13, 1030677.

34. Lim, I., Kim, B. C., Park, Y., Park, N. I., & Ha, J. (2022). Metabolic and developmental changes in germination process of mung bean (Vigna radiata (L.) r. wilczek) sprouts under different water spraying interval and duration. Journal of Food Quality, 2022(1), 6256310.

35. Maloney, G. S., Kochevenko, A., Tieman, D. M., Tohge, T., Krieger, U., Zamir, D., Taylor, M. G., Fernie, A. R., & Klee, H. J. (2010). Characterization of the branched-chain amino acid aminotransferase enzyme family in tomato. Plant Physiology, 153(3), 925–936.

36. Mengjun, L. G. (2019). TCseq: Time course sequencing data analysis. R package version, 1(0), 1–8.

37. Neinast, M., Murashige, D., & Arany, Z. (2019). Branched chain amino acids. Annual review of physiology, 81(1), 139–164.

38. Nonogaki, H. (2008). Seed germination and reserve mobilization. *eLS*.

39. Norton, L. E., & Layman, D. K. (2006). Leucine regulates translation initiation of protein synthesis in skeletal muscle after exercise. The Journal of nutrition, 136(2), 533S–537S.

40. Osmond, A. D., Directo, D. J., Elam, M. L., Juache, G., Kreipke, V. C., Saralegui, D. E., Wildman, R., Wong, M., & Jo, E. (2019). The effects of leucine-enriched branched-chain amino acid supplementation on recovery after high-intensity resistance exercise. International journal of sports physiology and performance, 14(8), 1081–1088.

41. Panchy, N., Lehti-Shiu, M., & Shiu, S.-H. (2016). Evolution of gene duplication in plants. Plant Physiology, 171(4), 2294–2316.

42. Pataczek, L., Zahir, Z. A., Ahmad, M., Rani, S., Nair, R., Schafleitner, R., Cadisch, G., & Hilger, T. (2018). Beans with benefits—the role of Mungbean (Vigna radiate) in a changing environment. American Journal of Plant Sciences, 9(7), 1577–1600.

43. Peng, C., Uygun, S., Shiu, S.-H., & Last, R. L. (2015). The impact of the branched-chain ketoacid dehydrogenase complex on amino acid homeostasis in Arabidopsis. Plant Physiology, 169(3), 1807–1820.

44. Platell, C., Kong, S. E., McCauley, R., & Hall, J. C. (2000). Branched-chain amino acids. Journal of gastroenterology and hepatology, 15(7), 706–717.

45. Robinson, M. D., McCarthy, D. J., & Smyth, G. K. (2010). edgeR: a Bioconductor package for differential expression analysis of digital gene expression data. Bioinformatics, 26(1), 139–140.

46. Rolletschek, H., Borisjuk, L., Gómez-Álvarez, E. M., & Pucciariello, C. (2025). Advances in seed hypoxia research. Plant Physiology, 197(1), kiae556.

47. Sá, A. G. A., Moreno, Y. M. F., & Carciofi, B. A. M. (2020). Plant proteins as high-quality nutritional source for human diet. Trends in Food Science & Technology, 97, 170–184.

48. Semba, R. D., Ramsing, R., Rahman, N., Kraemer, K., & Bloem, M. W. (2021). Legumes as a sustainable source of protein in human diets. Global Food Security, 28, 100520.

49. Shimomura, Y., Murakami, T., Nakai, N., Nagasaki, M., & Harris, R. A. (2004). Exercise promotes BCAA catabolism: effects of BCAA supplementation on skeletal muscle during exercise. The Journal of nutrition, 134(6), 1583S–1587S.

50. Sui, X., Zhang, T., & Jiang, L. (2021). Soy protein: Molecular structure revisited and recent advances in processing technologies. Annual Review of Food Science and Technology, 12(1), 119–147.

51. Tang, D., Dong, Y., Ren, H., Li, L., & He, C. (2014). A review of phytochemistry, metabolite changes, and medicinal uses of the common food mung bean and its sprouts (Vigna radiata). Chemistry Central Journal, 8(1), 4.

52. Taylor, N. L., Heazlewood, J. L., Day, D. A., & Millar, A. H. (2004). Lipoic acid-dependent oxidative catabolism of α-keto acids in mitochondria provides evidence for branched-chain amino acid catabolism in Arabidopsis. Plant Physiology, 134(2), 838–848.

53. Team, R. C. (2016). R: A language and environment for statistical computing. R Foundation for Statistical Computing, Vienna, Austria. http://www.R-project.org/.

54. Xing, A., & Last, R. L. (2017). A regulatory hierarchy of the Arabidopsis branched-chain amino acid metabolic network. The Plant Cell, 29(6), 1480–1499.

55. Yi-Shen, Z., Shuai, S., & FitzGerald, R. (2018). Mung bean proteins and peptides: Nutritional, functional and bioactive properties. Food & nutrition research, 62, 10.29219/fnr. v29262. 21290.

56. Yu, G., Wang, L.-G., Han, Y., & He, Q.-Y. (2012). clusterProfiler: an R package for comparing biological themes among gene clusters. Omics: a journal of integrative biology, 16(5), 284–287.

57. Zhao, M., Zhang, H., Yan, H., Qiu, L., & Baskin, C. C. (2018). Mobilization and role of starch, protein, and fat reserves during seed germination of six wild grassland species. Frontiers in Plant Science, 9, 234.

