## Supplementary Information for "Differential Regulation of Branched-Chain Amino Acids During Early Germination of Mungbean (*Vigna radiata* L.)"

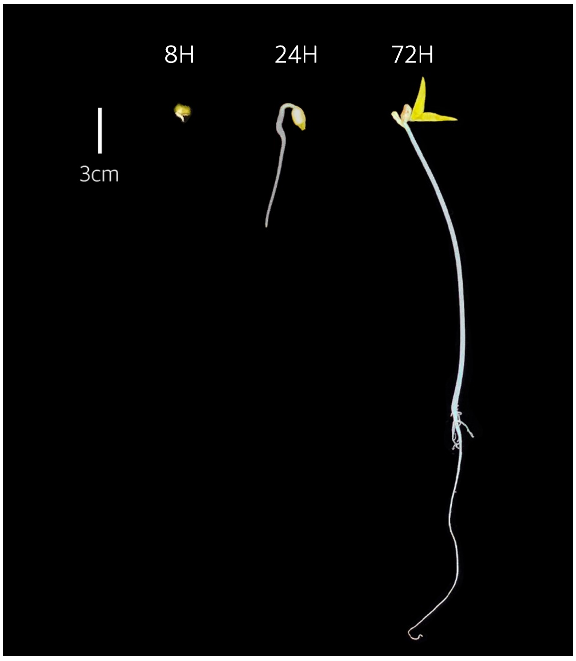


**Fig. S1**. Phenotypic appearance of mungbean sprouts during germination. Scale bar = 3 cm.

**Table S1**. Mobile phase gradient and flow rate in amino acid analysis.

| Time (min) | Pump 1 | | | | | | | Pump 2 | | | |
|---|---|---|---|---|---|---|---|---|---|---|---|
| | %B1 (PH-1) | %B2 (PH-2) | %B3 (PH-3) | %B4 (PH-4) | %B5 (PH-5) | Flow1 (mL/ min) | Temp (℃) | %R1 (reagent) | %R2 (buffer) | %R3 (water) | Flow2 (mL/ min) |
| 0.0 | 100 | 0 | 0 | 0 | 0 | 0.4 | 57 | 50 | 50 | 0 | 0.35 |
| 3.0 | 100 | 0 | 0 | 0 | 0 | | | | | | |
| 3.1 | 0 | 100 | 0 | 0 | 0 | | | | | | |
| 6.9 | 0 | 100 | 0 | 0 | 0 | | | | | | |
| 7.0 | 0 | 0 | 100 | 0 | 0 | | | | | | |
| 14.9 | 0 | 0 | 100 | 0 | 0 | | | | | | |
| 15.0 | 0 | 0 | 0 | 100 | 0 | | | | | | |
| 27.0 | 0 | 0 | 0 | 100 | 0 | | | | | | |
| 27.1 | 0 | 0 | 0 | 0 | 100 | | | | | | |
| 32.0 | | | | | | | | 50 | 50 | 0 | |
| 32.1 | | | | | | | | 0 | 0 | 100 | |
| 33.0 | 0 | 0 | 0 | 0 | 100 | | | | | | |
| 33.1 | 0 | 100 | 0 | 0 | 0 | | | | | | |
| 34.0 | 0 | 100 | 0 | 0 | 0 | | | | | | |
| 34.1 | 100 | 0 | 0 | 0 | 0 | | | | | | |
| 37.0 | | | | | | | | 0 | 0 | 100 | |
| 37.1 | | | | | | | | 50 | 50 | 0 | |
| 53.0 | 100 | 0 | 0 | 0 | 0 | | | | | | |

**Table S2**. RNA-seq quality metrics for mungbean sprouts at 8H, 24H, and 72H after germination. Metrics include total bases (bp), total reads, GC and AT content, base quality scores (Q20 and Q30), and alignment rate (%) for each replicate.

| Sample | Total bases (bp) | Total reads | GC (%) | AT (%) | Q20 (%) | Q30 (%) | Alignment rate (%) |
|---|---|---|---|---|---|---|---|
| 8H_1 | 5,981,435,938 | 59,222,138 | 45.1 | 54.9 | 97.7 | 93.6 | 97.3 |
| 8H_2 | 5,333,339,340 | 52,805,340 | 45.3 | 54.7 | 98.0 | 94.2 | 97.4 |
| 8H_3 | 7,694,222,218 | 76,180,418 | 45.0 | 55.0 | 98.0 | 94.3 | 97.5 |
| 24H_1 | 5,796,845,510 | 57,394,510 | 45.0 | 55.0 | 97.8 | 93.9 | 97.2 |
| 24H_2 | 7,701,258,282 | 76,250,082 | 44.8 | 55.2 | 97.8 | 93.9 | 97.3 |
| 24H_3 | 6,955,487,412 | 68,866,212 | 44.8 | 55.2 | 98.0 | 94.3 | 97.4 |
| 72H_1 | 6,373,359,570 | 63,102,570 | 45.1 | 54.9 | 97.9 | 94.2 | 97.3 |
| 72H_2 | 7,451,626,480 | 73,778,480 | 45.5 | 54.5 | 97.9 | 94.1 | 97.5 |
| 72H_3 | 5,102,853,300 | 50,523,300 | 44.9 | 55.1 | 97.9 | 94.0 | 97.3 |

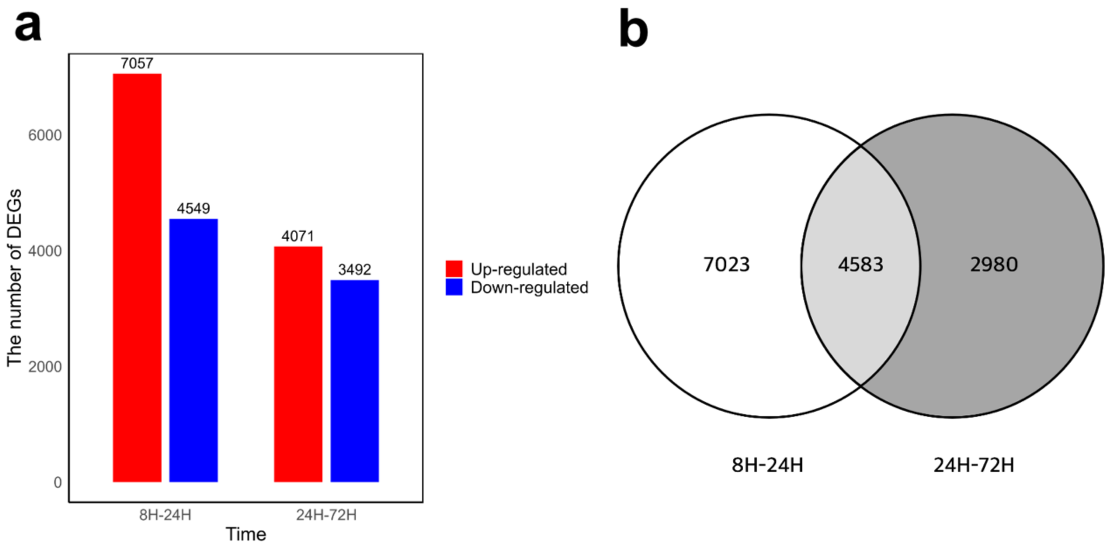


**Fig. S2**. Number of DEGs counts between each time point. (a) Number of DEGs identified between 8H–24H and 24H–72H during mungbean germination. Up-regulated (red) and down-regulated (blue) genes are shown separately. (b) Venn diagram showing the number of overlapping and unique DEGs between the two comparisons.

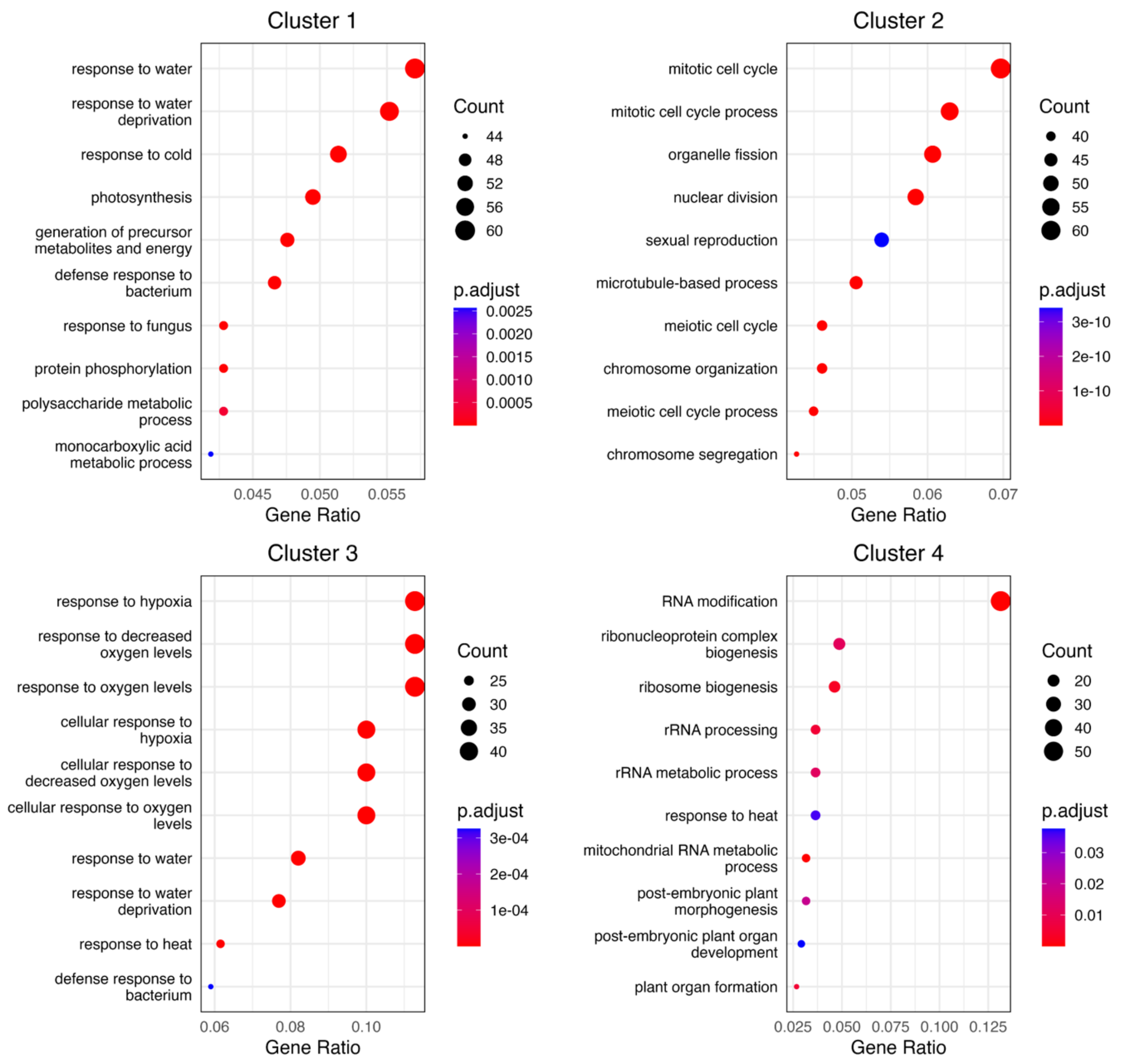


**Fig. S3.** GO biological process enrichment analysis of DEGs in each time-series expression cluster. GO biological process enrichment results for Clusters 1–4 identified in Figure 3. The size of each circle represents the number of genes, and color indicates adjusted *p* values (P.adjust).
